# Brain signatures for neuropsychological and everyday memory achieve high replicability and explanatory power in two data cohorts

**DOI:** 10.1101/2022.02.16.480746

**Authors:** Evan Fletcher, Sarah Farias, Charles DeCarli, Brandon Gavett, Keith Widaman, Fransia De Leon, Dan Mungas

## Abstract

**Background:** The “brain signature of cognition” concept has garnered interest as a data-driven, exploratory approach to better understand key brain regions involved in specific cognitive functions, with the potential to maximally characterize brain substrates of clinical outcomes. However, to be a robust brain phenotype, the signature approach requires a statistical foundation showing that model performance replicates across a variety of cohorts. Here, we outline a procedure that provides this foundation for a signature models of two memory-related behavioral domains.

**Method:** In each of two independent data cohorts, we derived regional brain gray matter thickness associations for neuropsychological and everyday cognition memory, testing for replicability. In each cohort we computed regional association to outcome in 40 randomly selected “discovery subsets” of size N = 400; we generated spatial overlap frequency maps and selected high-frequency regions as “consensus” signature masks for each cohort. We tested replicability by comparing cohort-based consensus model fits in all discovery sets. We tested explanatory power in each full cohort, compare signature model fits with competing “standard” models of each outcome.

**Result:** Spatial replications produced strongly convergent consensus signature regions derived from UCD and ADNI. Consensus model fits were highly correlated in 40 random subsets of each cohort indicating high replicability. In comparisons over each full cohort, signature models outperformed other models with one exception.

**Conclusion:** Multiple random model generations, followed by consensus selection of regional brain substrates, produced signature models that replicated model fits to outcome and outperformed other commonly used measures. Robust biomarkers of cognition and everyday function may be achievable by this method.

**Funding:** This project was funded by R01 AG052132 (NIH/NIA)

## Introduction

The “brain signature of cognition” concept has garnered interest as a data-driven, exploratory approach to better understanding key brain regions involved in specific cognitive functions, with the potential to maximally account for brain substrates of clinical outcomes. It has been characterized as discovering “statistical regions of interest” (sROIs or statROIs) (Chen et al., 2010; Fletcher et al., 2013; Hua et al., 2009) or brain “signature regions” associated with outcomes (Arenaza-Urquijo et al., 2019; Dickerson et al., 2009; Fletcher et al., 2021b; Gross et al., 2012). For a given brain measure, it computes areas of the brain that are most associated to a behavior outcome of interest. To be a robust brain phenotype, it requires validation, showing replicability in multiple datasets beyond those in which the signature was developed. Thus, signature models should be relatively independent of data cohort where generated and possess two key properties: 1) high and consistent explanation of outcome variance (i.e., model fit); and 2) consistent spatial location, leading to insights about brain substrates that underlie a behavior of interest.

In recent work (Fletcher et al., 2021b), we described a method for computing brain gray matter (GM) signatures of episodic memory in cognitively diverse populations and validated it across three independent cohorts. We found promising support for each of these properties. However, another recent study raised important questions about the replicability of exploratory approaches (Masouleh et al., 2019). Using multiple “discovery-confirmatory” sets to examine replicability of repeated signature generation and fit testing, they found little replicability for associations to cognitive outcomes in a cognitively healthy cohort, although distinct brain GM signatures of age, and to a lesser extent, BMI, were seen. In a second, clinical cohort that contained participants of varied cognitive status, they found better replicability for GM regions associated with immediate recall, both for model fits to behavior and consistent selection of memory-relevant brain areas by over 70% of the trials. Their results suggested that replicability of model fit and consistent spatial selection depend on cohort composition (particularly the need to capture a full range of variability in brain pathology and cognitive function), outcome domain of interest, and size of discovery set.

The present study thus has two aims. First, to develop and test a rigorous statistical foundation for the replicability and explanatory properties of the method laid out in our previous effort (Fletcher et al., 2021b). Second, to extend the method to another behavior domain in addition to neuropsychological measures of episodic memory: everyday memory function as measured by the Everyday Cognition scales (ECog), an informant-based scale for measuring subtle changes in day-to-day function of older participants (Farias et al., 2008). We investigate levels of attainable replicability both for model fits to outcomes and spatial selection of associated brain regions, and the generalizability of the signature approach to a second outcome domain.

## Methods

### Imaging cohorts

We used two imaging cohorts: 1) 578 participants from the UC Davis (UCD) Alzheimer’s Disease Research Center (ADRC) Longitudinal Diversity Cohort and 2) 831 participants from the Alzheimer’s Disease Neuroimaging Initiative phase 3 cohort (identified in the following as ADNI), downloaded from the ADNI site (adni.loni.usc.edu). All subjects had cognitive and everyday function (ECog) evaluations and one MRI scan taken near the time of evaluation.

One of the aims of the UCD ADRC cohort is to explore heterogeneity of cognitive trajectories in aging associated with a mixture of pathologies among an ethno-racially diverse group of older adults.

The ADNI project was launched as a public-private partnership in 2003 by the National Institutes of Aging, the National Institute of Biomedical Imaging and Bioengineering, the Food and Drug Administration, private pharmaceutical companies, and non-profit organizations. The primary goal of ADNI is to test whether serial MRI, PET, other biomarkers, and clinical and neuropathological assessment can be combined to measure progression of MCI and early Alzheimer’s Disease (AD). The principal investigator is Michael Weiner, MD, VA Medical Center and University of California, San Francisco. For current information on ADNI, see www.adni-info.org.

### Cognitive and everyday function assessment

Cognitive assessments of episodic memory were based on the Spanish and English Neuropsychological Assessment Scales (SENAS) (Mungas et al., 2005b, 2004) within the UCD ADRC cohort. SENAS is a composite measure based on a 15 item verbal list learning test incorporating performance across five learning trials and immediate recall. The memory composite from the ADNI cohort (ADNI-Mem) (Crane et al., 2012) was based on similar items from a list learning test as well as memory items from the Alzheimer’s Disease Assessment Scale-Cognitive Subscale (ADAS-Cog) and the Mini-Mental State Examination (MMSE). Both are sensitive to individual differences across the full range of episodic memory performance. The Everyday Memory domain from the ECog (ECogMem) (Farias et al., 2013, 2008) was used to measure everyday memory for both cohorts. The ECog is an informant-rated measure of several domains relevant to cognition as it applies to daily function. It was designed to address functional abilities of older adults, particularly focusing on subtle changes in everyday function spanning preclinical AD to moderate dementia (Farias et al., 2008).

### MRI image processing

We used single MRI scans in each cohort from the UCD and ADNI 3 cohorts. Whole head structural T1 MRI images were processed by in-house pipelines developed in our laboratory and described previously (Fletcher et al., 2014). The first pipeline step produced brain extractions based on convolutional neural net recognition of intracranial cavity followed by human quality control (Fletcher et al., 2021a). This was followed by affine and B-spline registration (Rueckert et al., 2006) of the intracranial cavity image to an age-appropriate structural template image (Kochunov et al., 2002) and native-space tissue segmentation into gray (GM) white (WM) and CSF (Fletcher et al., 2012) and white matter hyperintensities with the aid of each subject’s coregistered native T1 and FLAIR images (Decarli et al., 2013).

### Grey matter density quantification

We quantified brain cortical gray matter by gray matter density measures, performed at the voxel level in each native space image using the DiReCT diffeomorphic algorithm (Das et al., 2009) applied to the segmented gray matter. DiReCT is a ‘volume-based’ or 3D algorithm (i.e., it assigns a density measure to each GM voxel) as opposed the method employed in the commonly used Freesurfer package, which is ‘surface-based’ (calculating vertex-wise distances between inner and outer 2D GM surface meshes) (Fischl and Dale, 2000). Resulting native-space GM density maps were deformed to template space using the affine and B-spline parameters previously computed in our pipeline.

### Signature variable analyses

We computed signature models in each cohort. An algorithm overview appears in Fig. 1. In Step 1, we generated signature masks of GM thickness association to outcome in each of 40 randomly selected discovery sets (N = 400 for each discovery set) within each cohort. Separate masks were generated at each of three levels of association using regression β coefficient t-values (t = 3, 5, 7). In step 2, we combined all 40 signature masks into cohort-specific overlap frequency maps, then selected cohort “consensus” signature masks consisting of overlap frequency greater than 70 percent at each of three t-levels of association. In the validation steps, we tested cohort consensus signature models by comparing their performances with each other in each of the discovery sets of each cohort. Finally, we also compared consensus signature models with other, competing models of outcome in the full UCD and ADNI cohorts.

**Figure 1.**
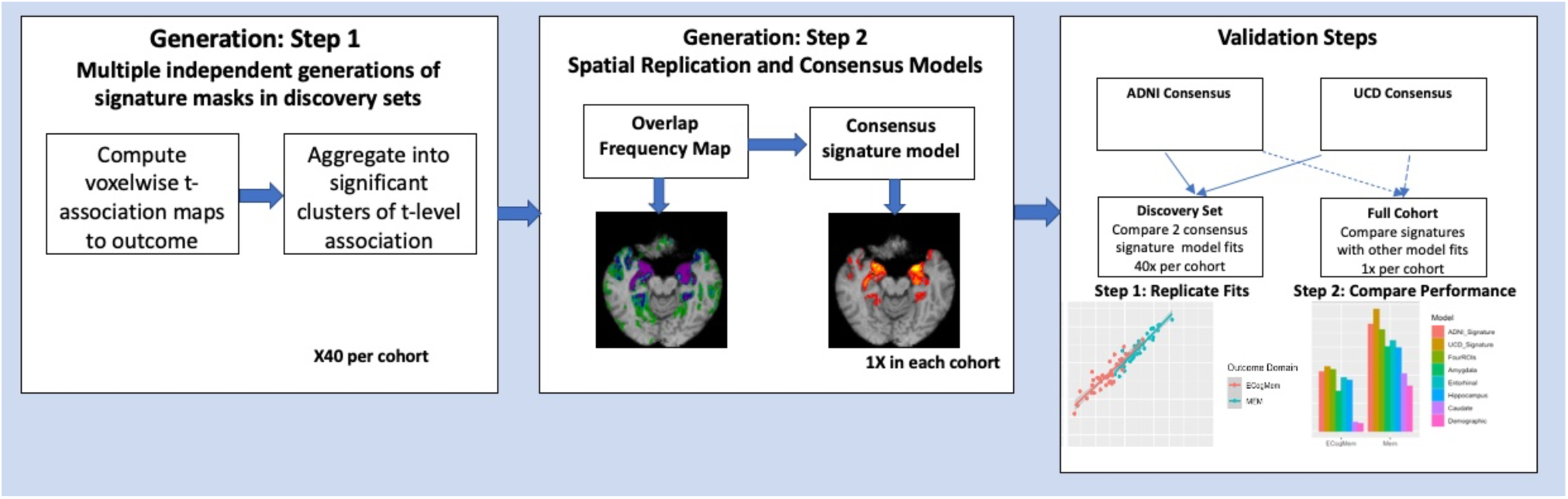
Overview of analysis steps and sample outputs at each step.

#### Generation Step 1. Signature mask computation

This step is summarized in Fig. 1, left box. Details are shown in Fig. 2A, B. We generated signature masks of association to outcome domain independently in each discovery set. This process was extensively described in our previous work (Fletcher et al., 2021b) and will be briefly summarized here. Regressions were performed at each template-space voxel with outcome domain as the dependent variable, GM density as the independent variable of interest, and controlling for age, gender, and education. The resulting GM maps of voxel-based t-values (i.e., the t-value of regression β coefficient for GM density) indexed the strength of association of GM at every voxel. To ascertain statistical significance while aggregating the voxels into significant clusters, we performed non-parametric t-value cluster size computations (Nichols and Holmes, 2001) using 10,000 iterations separately for t-thresholds of 3, 5 and 7. This computed an empirical distribution of cluster sizes under the null hypothesis of no association between brain and behavior outcome. Clusters from the original regressions with size in the top 5% (95^th^ percentile) of this distribution were retained as significant. In practice, most regions of interest selected for signature masks were in the highest 0.01% (i.e., they were the largest clusters over the 10,000 repetitions). Each discovery set thus produced three significant clusters, corresponding to the three t-values, for each outcome domain. These were the signature masks for a discovery set, designated as TsROIi for i = 1, 2, 3, corresponding to t-values of 3,5,7.

**Figure 2.**
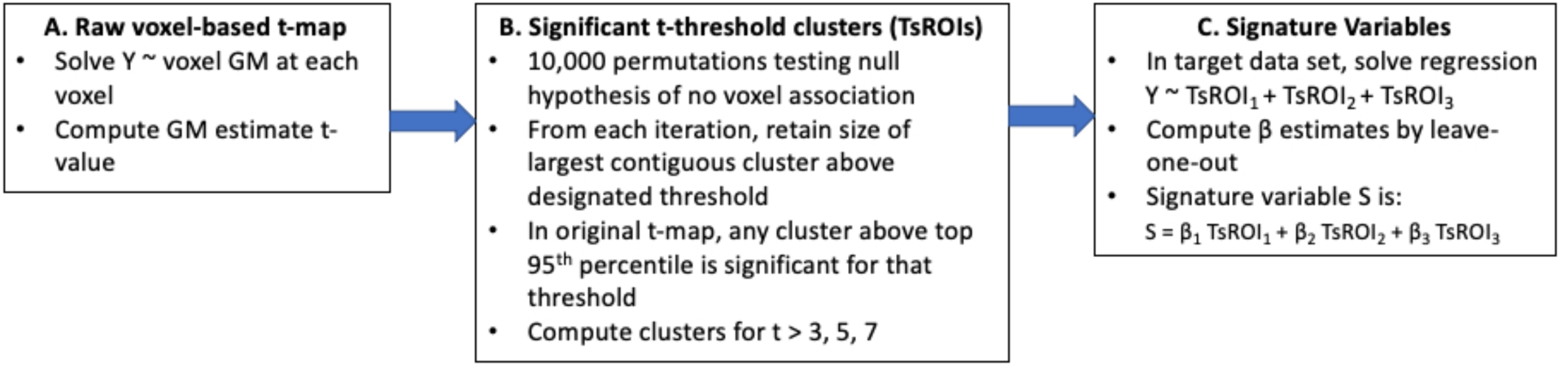
Steps to create signature variables. Steps A and B are performed in the discovery set to learn the TsROI masks. Step C is performed in a target set where signature variables are required.

#### Signature variable computation

This step is shown in Fig. 2C. In a dataset of interest (the target dataset), we sampled mean GM cortical density in each of the TsROIs for each subject. Signature variables in the target set are linear combinations of the TsROI variables computed by solving the regression equation (1) in a leave-one-out manner.

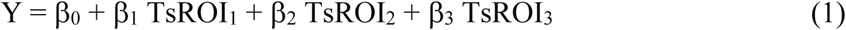

Here Y is our selected behavior outcome and the TsROIs are corresponding GM density means for each subject. Equation (1) allows the β coefficients to vary by target set. This increases the explanatory power of the TsROI models (Fletcher et al., 2021b). However, to minimize possible target set-related bias in β computations, we employed a leave-one-out procedure in the target set. Thus, each subject was held out in turn and the β coefficients were computed in a regression using the remaining N-1 subjects. The left-out subject’s predicted outcome (its signature variable S) was obtained by inserting the computed β values and the left-out subject’s TsROI values in the right side of equation (1).

#### Generation Step 2. Consensus signatures

This step is shown in Fig. 1, middle box, with details in Fig. 2C. We computed overlap frequency maps of TsROI masks from all 40 discovery sets. We then defined our consensus signature masks at each level of t = 3,5,7 as the 70% or higher overlap regions in our frequency maps for a given t-level. To compute consensus signature variables for an outcome in a target set, we used the three consensus TsROI masks with GM means and the β coefficients computed in the target set.

#### Testing replicability and validating signature models

These steps are summarized in Fig. 1, right box. In each step we utilize consensus signatures generated in both UCD and ADNI cohorts.

#### Validation Step 1. Testing replicability for two cohorts of origin

In each of 40 discovery sets in a cohort, we computed signature variables from consensus TsROI masks generated in that cohort and in the opposite one. We then computed the fit of each signature variable S to outcome, controlling for age, gender, and education (equation (2)).

Overall fit was measured by adjusted R^2^.

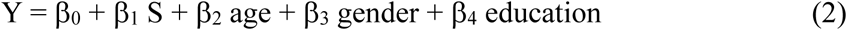

For comparison, we also computed explanation of variance by demographics alone:

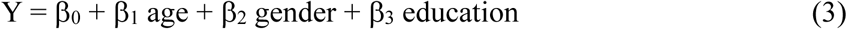

To estimate the incremental variance ΔR^2^ explained by S beyond demographics, we subtracted the fit coefficient of equation (3) from the fit of the full model (2): Δ R^2^ = adjusted R^2^ (full) – adjusted R^2^ (demographic).

#### Validation Step 2. Testing optimal performance

Our final test compared the fit performances of the consensus signatures against those of other brain variables within each of the entire ADNI and UCD cohorts. From a cortical parcellation atlas (https://mindboggle.info/) we selected four regions most heavily overlapped by each of the consensus masks in at least one cohort. These were the amygdala, entorhinal cortex, hippocampus and caudate. We regressed an outcome on each of these variables in turn, controlling for age, gender, and education, and tabulated the adjusted R^2^ fit measures (not the ΔR^2^). In addition to these single-ROI models, we made a multivariate model incorporating all these ROIs model as predictors. And our last comparison examined the outcome adjusted R^2^ for the three demographic variables alone.

## Results

### Cohort Demographic Profiles

Participant and scanner characteristics of our two cohorts are presented in Table 1. The ADNI cohort was significantly younger (p < 0.001), significantly less female (p < 0.001) and had significantly more education (p < 0.001) than UCD. ADNI was almost entirely non-Hispanic / Latino whereas UCD had about 50% white and almost one-quarter each of African American and Hispanic / Latino. For clinical diagnosis, UCD had a significantly greater proportion of normal (CN) than ADNI (p = 0.008 via likelihood ratio) as well as significantly more participants with dementia (p < 0.001). In ADNI, the clinical diagnoses are principally in the Alzheimer’s spectrum, whereas the UCD demented category included vascular as well as Alzheimer’s disease dementias. Our last measure of scanner field strengths shows that ADNI consisted of almost entirely 3T scanners (in fact there was just one 1.5T) whereas UCD was about 78% 1.5T.

**Table 1.** Demographic profiles of the two cohorts. Abbreviations: CN = cognitively normal; MCI = mild cognitive impairment; AD = Alzheimer’s disease

|  | **ADNI 3** | **UCD** |
| --- | --- | --- |
| **n** | 815 | 576 |
| **Age, years (mean (SD))** | 71.4 (7.3) | 76.8 (7.0) |
| **Gender (percent female)** | 52 | 60 |
| **Education, years (mean (SD))** | 16.5 (2.5) | 13.9 (4.1) |
| **Race/ethnicity (percent)** | Hispanic/Latino 4<br>Not Hispanic/Latino 95 | Asian 3<br>African American 23<br>Hispanic/Latino 21<br>White 50<br>Other 2 |
| **Clinical Diagnosis (percent)** | CN 50<br>MCI 27<br>AD 10<br>Not available 13 | CN 56<br>MCI 26<br>Demented 17 |
| **Scanner Field Strength (Percent 3 Tesla)** | 99.9 | 22 |

### Replication of spatial selection: overlap frequency maps

Fig. 3 displays overlap frequency maps for cognitive memory-association clusters in each cohort. ECog memory (not shown) exhibited similar patterns, with one exception: the caudate was not selected by UCD ECog signatures. Maps in each cohort show strong consensus overlaps (purple: 100%) for medial temporal, amygdala, and hippocampal locations. More dorsally, there was strong overlap in the caudate, though with somewhat smaller extent of the 100% regions than in the temporal slices. We also note small areas of high-frequency overlaps in the precuneus and PCC for both ADNI and UCD (rightmost slices in each cohort).

**Figure 3.**
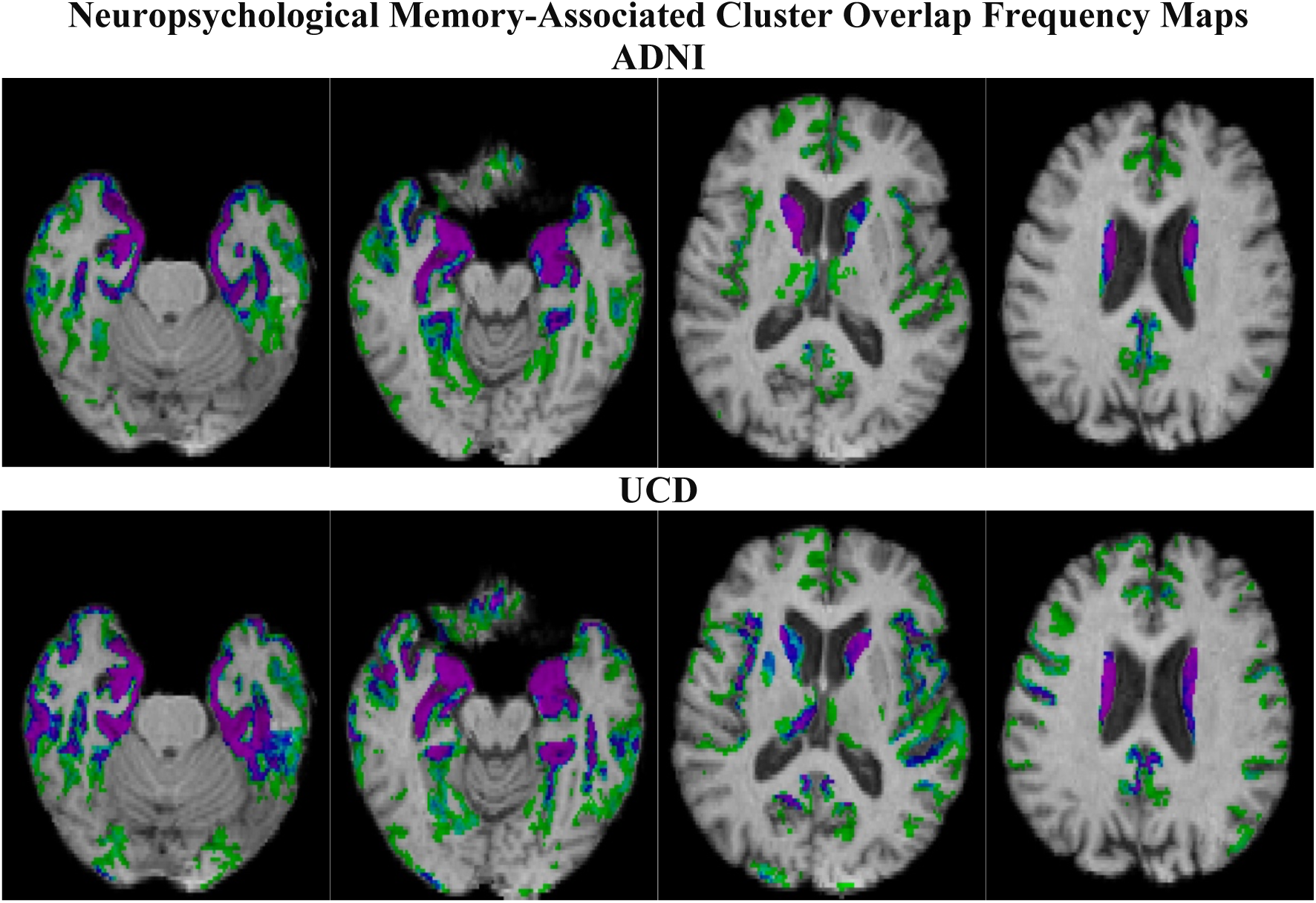
Percentage overlap of significant gray matter cluster associations at t = 3 to memory over 40 random trials in each cohort. Percentage frequency coding goes from light green (2.5%, i.e., 1/40) to purple (100%, i.e., 40/40)

### Cohort-generated consensus signature models

Fig. 4 displays the color-coded consensus TsROI signature maps for ADNI and UCD, Memory and ECog Mem. These images closely follow the high-frequency consensus levels (blue to purple) in Fig. 3, but now regions of variable t-strength association are also visible.

**Figure 4.**
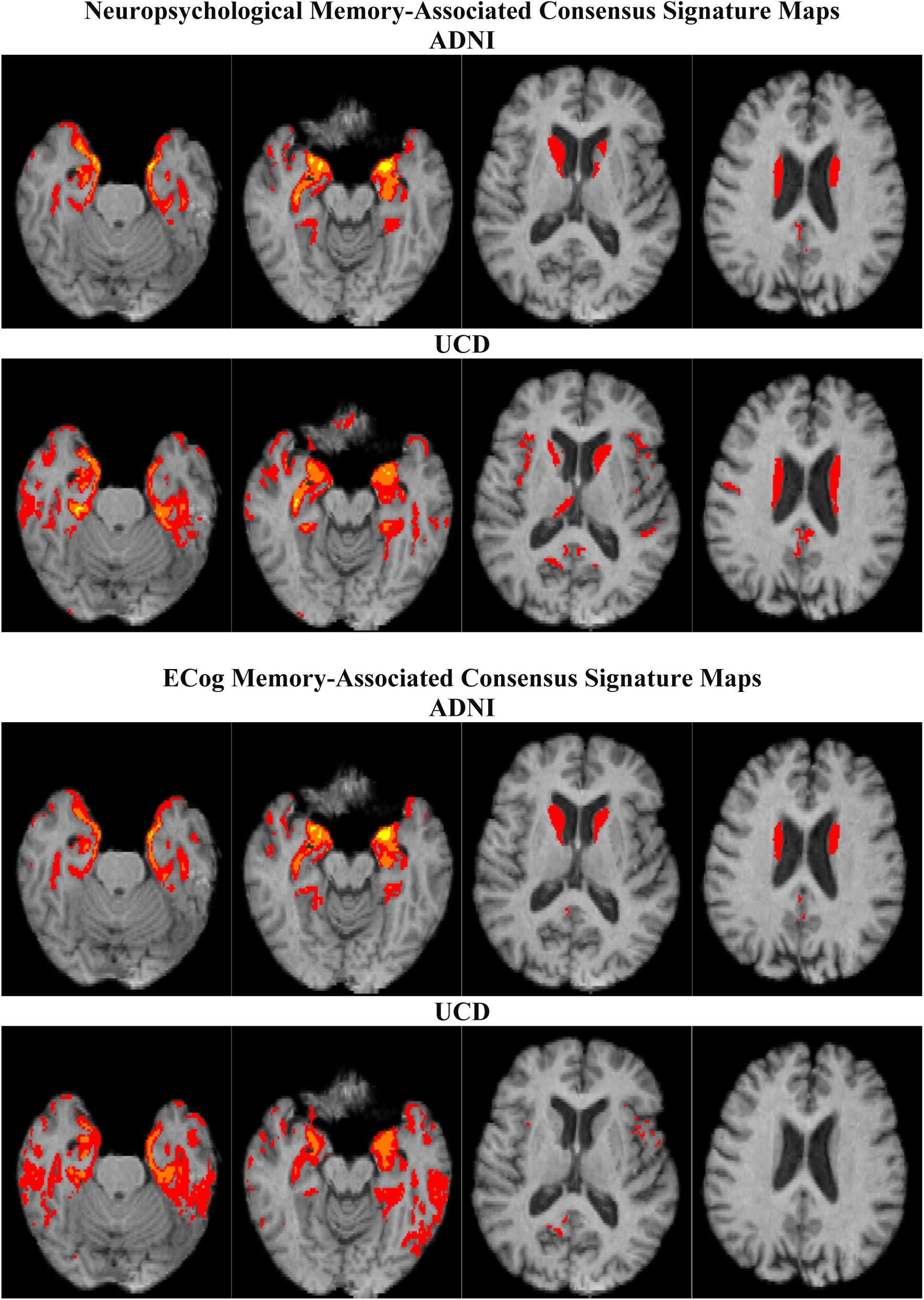
Consensus signature TsROI regions for memory (top) and ECog Mem (bottom) computed in each cohort. Based on 70% overlap “consensus” at each of three t-levels of association: t = 3 (red), 5 (orange), 7 (yellow).

For neuropsychological memory, consensus masks in ADNI and UCD show a strong convergence, each cohort having extensive associations at levels t = 3, 5 and 7 within temporal regions (leftmost images), and associations at t = 3 within the caudate and the posterior cingulate / precuneus (rightmost images). For ECog Memory, the cohort consensus TsROIs are also convergent in the two cohorts, except that the UCD TsROIs do not show any association of caudate with ECog outcome. We also note that in each cohort, the signature TsROIs for neuropsychological Memory and ECog Memory are similar, but again the UCD ECog Mem signature differs from its Memory signature by the absence of overlap in the caudate. In sum, consensus signature masks show strong resemblances for ADNI vs. UCD by outcome domain, and strong similarities between outcome domains by cohort of origin.

Table 2 shows the percent overlap of selected brain atlas regions by consensus signature masks (t >= 3), in other words by all the colored regions displayed in Fig. 4. All four signature maps overlapped three medial temporal structures (amygdala, entorhinal cortex, and hippocampus), at consistently high percentages of those structures (roughly 60-95%) and except for the two UCD ECog signatures, around 45% of the caudate. The parahippocampal gyrus was overlapped at mid 40% levels by the UCD signatures but also consistently at lower percentages by the ADNI signatures. Similar patterns are seen for the fusiform and inferior temporal regions.

**Table 2.** Top 15 regional atlas overlaps for consensus masks corresponding to t >= 3, sorted for the UCD memory signature overlaps.

|  | ADNI 3 |  | UCD |  |
| --- | --- | --- | --- | --- |
|  | ECogMem | Memory | ECogMem | Memory |
| Amygdala | 0.94 | 0.93 | 0.91 | 0.98 |
| Entorhinal | 0.72 | 0.74 | 0.78 | 0.77 |
| Hippocampus | 0.59 | 0.63 | 0.65 | 0.64 |
| Parahippocampal | 0.34 | 0.29 | 0.46 | 0.46 |
| Caudate | 0.47 | 0.43 | 0.03 | 0.44 |
| Isthmus Cingulate | 0.19 | 0.12 | 0.12 | 0.29 |
| Fusiform | 0.14 | 0.13 | 0.31 | 0.26 |
| Inferior Temporal | 0.10 | 0.07 | 0.49 | 0.22 |
| Medial Orbitofrontal | 0.09 | 0.05 | 0.13 | 0.21 |
| Insula | 0.04 | 0.07 | 0.16 | 0.19 |
| Pars Orbitalis | 0.0 | 0.0 | 0.06 | 0.18 |
| Superior Temporal | 0.15 | 0.11 | 0.17 | 0.13 |
| Lateral Orbitofrontal | 0.02 | 0.01 | 0.10 | 0.13 |
| Transverse Temporal | 0.0 | 0.0 | 0.0 | 0.13 |
| Middle Temporal | 0.03 | 0.02 | 0.21 | 0.12 |

### Cross-cohort performance replications of consensus signature variables

We next examined the comparative performances of same-and opposite-cohort consensus model fits for each domain in the 40 discovery sets of each cohort. Here, “same-cohort” consensus signatures used consensus TsROIs derived in the same cohort as the target set, and “cross-cohort” signatures came from masks generated in the other cohort. Thus, in each of the discovery sets for each cohort, we computed the signature model fit, controlling for age, gender, and education, of each cohort consensus signature with its appropriate outcome domain (equation (2)), and then the ΔR^2^ for variance explained beyond demographic variables.

The results are displayed in Fig. 5. The top panel shows scatterplots for adjusted ΔR^2^ fits. There are very tight correlations across 40 trials in each cohort. The bottom panel shows density plots of the ratios of the ΔR^2^ values. For cognitive memory, the consensus variables achieved very tight peaks with the cross-cohort ΔR^2^ being approximately 0.8 times the within-cohort consensus fit. This is seen directly in the ADNI panel, where UCD is the cross-cohort variable, and inversely in UCD, where ADNI is the cross-cohort variable. ECogMem is also sharp in ADNI and less so in UCD. In sum, overall model fits for the consensus signature variables are highly correlated regardless of cohort of origin or target cohort where evaluated, and the cross-cohort signature fit metric is tightly clustered at about 0.8 times the within-cohort metric.

**Figure 5.**
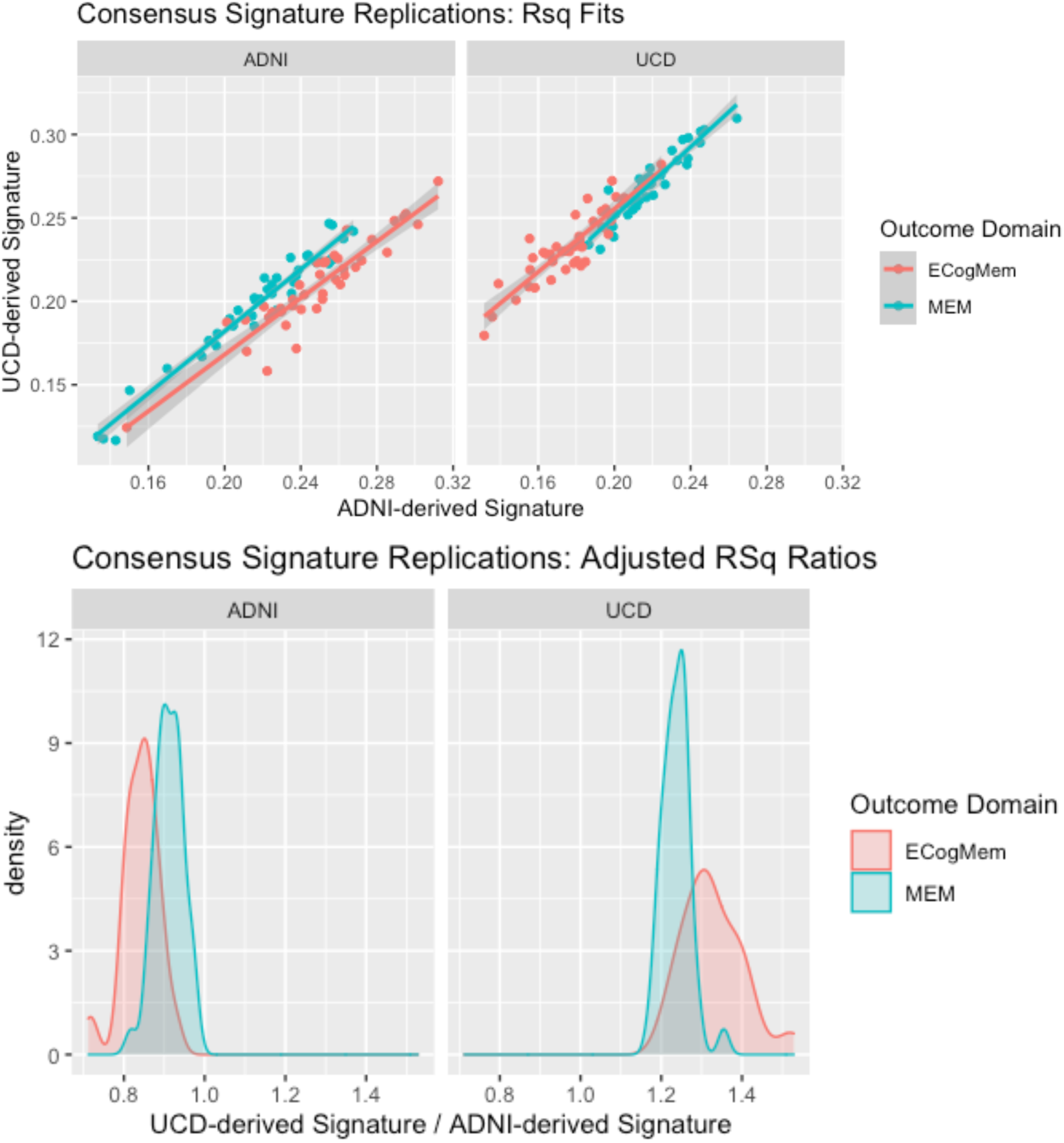
Plots of cohort consensus signature model fits for ΔR^2^ (incremental adjusted R^2^ above demographics alone) in the 40 discovery sets of each trial cohort. Fits for outcome domain are regressed on signature variables, controlling for age, education, and gender. Top: scatterplots for y = ΔR^2^ of UCD consensus models, x = ΔR^2^ of ADNI consensus models. Bottom: density plots of the ratios of these ΔR^2^ fits. Mirror symmetry is because UCD was the cross-cohort consensus signature in ADNI but the same-cohort signatures in UCD.

### Optimal performance of consensus signature models

Our final test examined the fit performances of each cohort consensus model in each full cohort. Fits were measured from adjusted R^2^ of regressions including age, gender, and education as covariates (equation (2)). We did not compute ΔR^2^ because we were interested in the comparative performances of full models, including demographics that were included as covariates. We compared signature model performances to those of brain regions figuring prominently in the consensus models: entorhinal cortex, amygdala, hippocampus and caudate, and finally a model incorporating all four of these regions as predictors (FourROIs). To demonstrate a baseline level of predicted variance from demographic factors alone, we included fits for the model incorporating age, gender, and education but no brain predictors. Results are displayed in Fig. 6.

**Figure 6.**
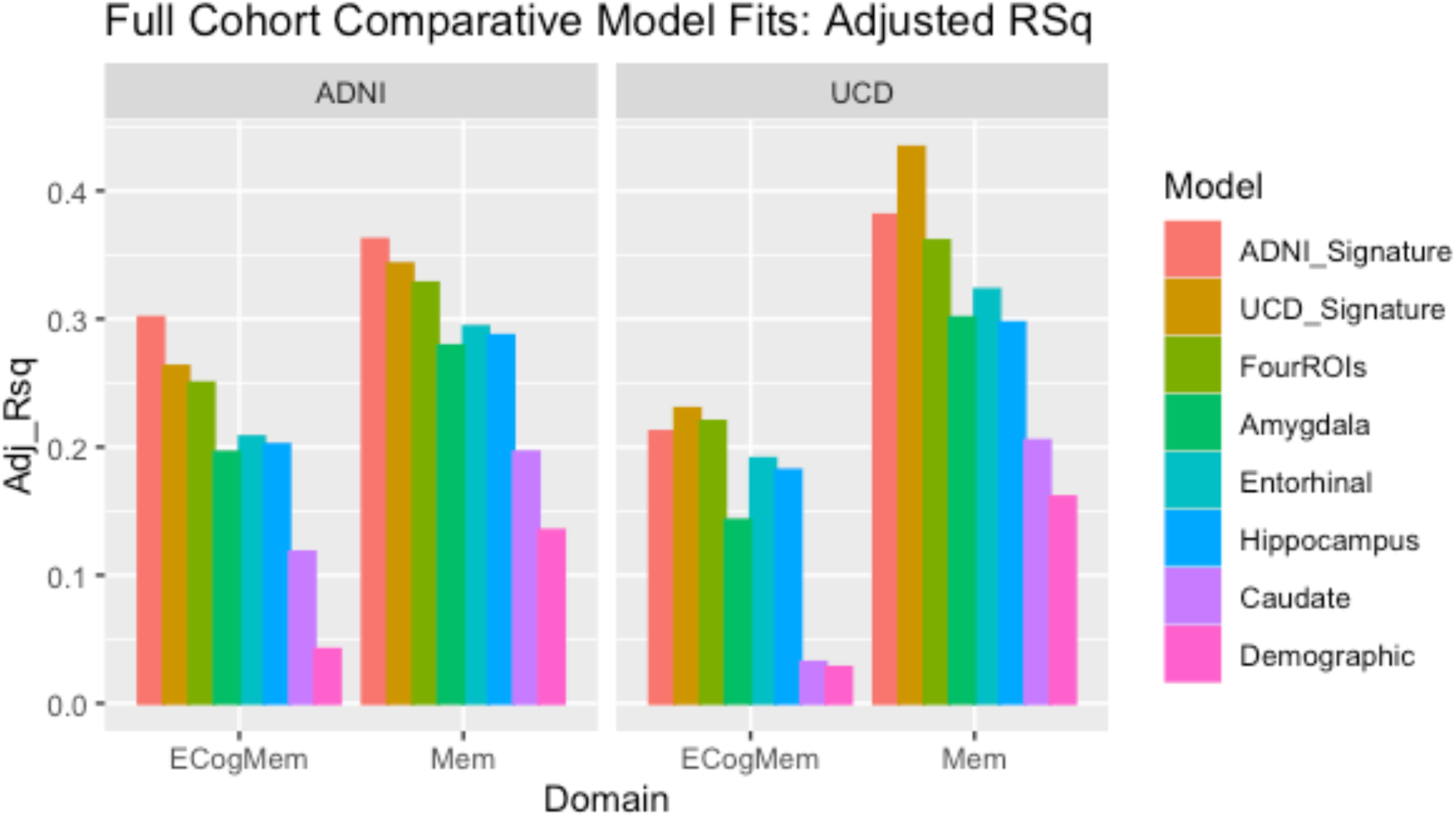
Comparative model fits (measured by adjusted R^2^ in regressions controlling for age, gender, and education) in the entire UCD and ADNI cohorts. FourROIs designates the model incorporating all of amygdala, entorhinal, hippocampus and caudate as multivariate predictors. Demographic refers to a model incorporating age, gender and education without any brain predictors. All models incorporate demographic variables, and demographic has no brain variables.

As expected, each cohort consensus model performed the best within its own cohort, outperforming the signature model of the opposite cohort as well as all competing single-region models. The second-highest fit was invariably the consensus model of the other cohort, except for ECogMem in UCD, where the FourROIs model was slightly higher than the ADNI Signature. The Demographic models showed lowest fits among all models, demonstrating that brain factors add improved outcome variance prediction above demographic factors alone. Interestingly, the demographic models explained less of ECogMem variance than of Memory, thus perhaps accounting for the lower fits of the ECogMem consensus models that controlled for demographic variables. The FourROIs models showed higher fit than any individual brain component but were still exceeded by both same-cohort and cross-cohort consensus signature models, with the one exception (barely) just noted. In sum, the consensus signature variables attained optimal or near-optimal fit performance among the competing models tested.

## Discussion

### Summary of method and results

This project had two aims. First, we aimed to develop a rigorous statistical foundation, based on tests of replicability, of the exploratory voxel-based signature approach documented in our recent publication (Fletcher et al., 2021b). Second, we aimed to extend the exploratory approach beyond neuropsychological memory to a second outcome domain of everyday cognition (ECogMem) (Farias et al., 2013, 2008), investigating similarities in brain GM substrates for these outcomes.

Our approach used multiple trials of signature mask generation in each of two imaging cohorts, leading to maps of spatial overlap frequencies from which we selected consensus regions having high spatial selection frequency across all trials. We then tested model fit replicability of consensus signature models generated in each of the cohorts. We performed these steps using the recommendations of (Masouleh et al., 2019) for large signature discovery sets drawn from cognitively heterogeneous cohorts.

### Spatial and model fit replicability

In each of two cognitively heterogeneous cohorts, 40 independent computations of ROIs associated to outcome showed high spatial replicability (Fig. 3), allowing us to designate consensus regions by cohort (Fig. 4). Signature variables computed from cohort consensus regions achieved model fits of outcome that were highly correlated across 40 randomly chosen sets in each cohort (Fig. 5). Thus, leveraging spatial replicability across multiple trials led to consensus signature variables having high model fit replicability, validating these signature regions as useful brain biomarkers. Finally, we found that these signature models achieved better explanations of outcome variance than other plausible and standardly used models (Fig. 6). To the best of our knowledge, the algorithm producing rigorous validation and high fit performance is a novel contribution of this work.

### Brain GM substrates of neuropsychological and everyday memory

We note strong similarities of neuropsychological memory and ECog memory consensus brain signatures (Fig. 4), with one exception of missing caudate for the UCD ECog signature. Thus, convergent signature spatial locations suggest shared brain GM substrates including medial temporal regions for both memory outcomes (Fig. 4), and lesser but consistent overlaps with isthmus cingulate (Table 2). All these regions are known to be involved in episodic memory. They are structurally and functionally connected (Fjell et al., 2016, 2015) and are part of the default mode network (DMN) that is vulnerable to Alzheimer’s disease (Greicius et al., 2009). Neuropsychological memory signatures also overlap the caudate, while the absence of this structure in the UCD ECogMem signatures warrants further investigation into cohort-based differences of factors known to moderate associations of brain substrates to outcome, e.g. (Gavett et al., 2018).

Beyond the difference of caudate overlap, overall signature similarities exist between neuropsychological and ECog memory, despite the fact that neuropsychological memory (Mungas et al., 2005a, 2004) and ECogMem (Farias et al., 2008) are evaluated by different metrics (in particular, the version of ECog used here relies on third-person informant reports). Earlier research found correlations of ECog memory with brain measures (hippocampal and total brain volume, dorsolateral prefrontal cortex) and neuropsychological memory (Farias et al., 2013). Our findings for signature regions of ECog memory are thus consistent with the previously found regional associations, extending them to a robust similarity of overall brain GM substrates for both memory-based outcomes.

Finally, looking at the rough equality of ΔR^2^ magnitudes for neuropsychological memory and ECogMem (Fig. 5) suggests that the amount of outcome variance explained *by the signatures* is similar for both. In sum, it may be that neuropsychological memory and ECogMem share similar brain GM substrates, with about the same strengths of association to these substrates. This is a potentially new finding and a topic for further research.

### Relations to previous signature models

In the signature models of our previous study (Fletcher et al., 2021b), validation was confined to observations of model fits in each of three target cohorts. The present work extends that study by showing consistency and replicability over 40 randomly selected sets in two independent imaging cohorts, and over those full cohorts as well. Comparative whole-cohort model fits displayed in Fig. 6 of the current study are in line with those found in our previous work, although our current adjusted R^2^ are somewhat higher overall, probably because of generally larger discovery data sets in the current study. This suggests that the consensus and leave-one-out steps introduced here did not degrade model fit performance compared to our earlier work, and the consensus procedure may have enhanced it, while also providing better verification of replicability.

The recent empirical examination of signature replicability (Masouleh et al., 2019) suggested the multiple trials approach we followed here. They reported little replicable association between brain and behavior in a cognitively normal cohort, but found some regions selected by more than 70% of the trials for brain GM associations with short term memory in a cognitively mixed, clinical cohort. Reassuringly, many of their regions in the clinical cohort appear similar to our consensus signature TsROIs. We thus may have corroborated their results for a cognitively mixed cohort, and with even stronger associations, perhaps due to our use of larger discovery set sizes, consistent with the recommendations of that work.

### Relations to brain atlases and theory-driven models

High quality brain image parcellation atlases, e.g., (Klein et al., 2017; Manera et al., 2020) have many benefits. They may be used directly in arbitrary study cohorts to implement “theory-driven” models based on accumulated findings on the relationship of brain structures to behavior. They do not require exhaustive exploratory search procedures or verification of replicability.

They constitute a form of “data reduction” that is valuable for constructing tractable models. On the other hand, atlas regions do not necessarily align with networked locations that underlie behavioral outcomes of interest (Jolly and Hampshire, 2021) and this may explain why their explanatory performance falls short of that attained by signature models, seen in this work (Fig. 6) and our previous study. Recent efforts have incorporated both atlas and exploratory concepts by searching lists of parcellation ROIs for a subset that optimally explains an outcome of interest (Epelbaum et al., 2018; Schwarz et al., 2016). However, this procedure may ignore the partial rather than full involvement of selected ROI locations, and as our previous publication showed, their fit performance was below that of our signature region models. The FourROIs model from the current study illustrates this. Though it incorporated the four atlas regions most heavily overlapped by our signature masks, its fit performance was still lower than the signatures except in one case. Thus, one of the advantages of the exploratory signature approach may be that its selection process implicitly accounts for connections of disparate regions communicating with each other in networks associated with behavioral outcomes (Genon et al., 2018; Jolly and Hampshire, 2021). This may contribute to its superior explanatory power over that of pre-selected regions.

### Strengths and Limitations

An important strength of the signature approach is that it proposes hypothesis-free, exploratory computation of brain regional measures maximally associated to outcome (Bakkour et al., 2009; Dickerson et al., 2009; Fletcher et al., 2021b; Jolly and Hampshire, 2021). However, achieving this promise necessarily incurs substantial difficulties (Masouleh et al., 2019). First, a definitive brain signature of an outcome may not even exist, due to individual differences and the lack of specificity in brain-behavior relations (Genon et al., 2018). Second, poor replicability and even lack of association between brain and outcome in a population may challenge the validity of the signature concept (Masouleh et al., 2019). Overcoming this challenge requires the availability of large and varied data sets to obtain robustness. Third, behavioral outcomes depend on arrays of factors that are difficult to fully account for (Habes et al., 2020). An appropriate implementation of the signature approach may come from machine learning, which is capable of accounting for interactions between many more factors than human-constructed models could incorporate (Dinsdale et al., 2020). But machine learning entails its own challenges, including the need for very large data sets (Fletcher et al., 2021a) and an opacity of output that may not be readily interpretable or accessible to human understanding (Böhle et al., 2019).

One aim of our current effort has been to address the replicability issues raised by (Masouleh et al., 2019). By using multiple iterations of large discovery sets in cognitively heterogeneous cohorts, we have improved on the replicability they reported for the clinical cohort they used.However, our discovery set trials, like theirs, were not truly independent since the randomly chosen discovery sets necessarily overlapped. And in any case, there remain differences by cohort in brain configurations of our signatures. A next step may use even larger data sets with harmonized outcomes to see if these can be reduced. On the other hand, to the extent that our results provide confidence of robust replicability, these differences may themselves suggest fruitful lines of further research aimed at more clearly delineating brain-behavior relations and how they are modified by other non-signature variables in diverse populations.

## Conclusion

In conclusion, our results support the feasibility of generating behavior-related brain signatures that depend minimally on discovery set of origin and can be used as robust GM brain phenotypes. This is one novel finding of our work. For behavior outcomes of memory and everyday function, we found that the associated brain phenotypes are strongly convergent. This is another new finding that warrants further exploration. Lines of future research could include using larger datasets for exploring cohort-based differences in signature models, limits of replicability, and developing signature models accounting for an array of brain measures beyond GM.

## Notes

http://adni.loni.usc.edu/

### Competing Interest Statement

The authors have declared no competing interest.

